# Coastal management challenges in declining clam fisheries: insights from local ecological knowledge in Spain

**DOI:** 10.64898/2026.02.08.704719

**Authors:** Marc Baeta, Laura Benestan, Marco Antonio Solis, Mauricio Mardones, Marina Delgado, Luis Silva, Miguel Rodilla, Silvia Falco, Manuel Ballesteros, Miriam Hampel, Ciro Rico

## Abstract

Spanish clam fisheries have experienced declining landings and repeated closures, raising questions about how ecological pressures and governance constraints are understood by those involved in their management. We conducted 94 semi-structured interviews with fishers, fishers’ guild leaders and regional managers across Spain’s Mediterranean and southern Atlantic coasts between April 2024 and August 2025. Focusing on (*Donax trunculus*), (*Chamelea gallina*) and (*Callista chione*), we used thematic analysis to examine perceived pressures, institutional constraints and contextual variation among study areas. Interviewees identified multiple pressures, including environmental variability, habitat alteration, pollution, episodic mortality and fishing-related factors. Assessments of management and governance were predominantly negative, with recurring concerns about coordination, participation, uneven implementation of rules, monitoring and enforcement. Accounts also highlighted conflicts with other coastal users and differences in the fishing contexts represented across study areas. These findings are consistent with a context-dependent understanding of decline in which perceived ecological pressures and institutional constraints are closely connected. However, the purposive sample does not support population-level estimates, causal attribution, or independent validation of management effectiveness. Policy options arising from the analysis include more coordinated rules across modalities, participatory monitoring, traceability and enforcement adapted to nearshore conditions, and stronger integration of fisheries into coastal decision-making. The study identifies priorities for more coherent governance while distinguishing stakeholder accounts from demonstrated ecological mechanisms and management outcomes.

**Highlights:**

- Stakeholders identify multiple pressures associated with clam decline.
- Perceived drivers and management concerns vary among coastal settings.
- Interviewees report coordination, participation and enforcement constraints.
- Shore-based fisheries present distinctive monitoring and access challenges.
- The findings inform options for more integrated coastal governance.

## 1. Introduction

Coastal fisheries increasingly operate in crowded, rapidly changing nearshore environments where ecological dynamics, market conditions, and governance arrangements interact. For low-mobility resources such as clams, these interactions can translate into prolonged declines that are difficult to reverse once recruitment, habitat suitability, or compliance capacity are compromised. This pattern has been widely recognised in policy discussions on the vulnerability of small-scale fisheries (SSF) to environmental variability and socio-economic shocks, and on the need for governance systems that are both legitimate and adaptive (FAO, 2024; Jentoft, 2007).

Spanish clam fisheries, as a form of small-scale fisheries (SSF), provide a policy relevant case of this coupled social-ecological challenge. Like other coastal SSF, they are strongly place based, economically important for some coastal communities, and embedded in nearshore spaces where fishing activities interact with other activities and competing uses (Bennett et al., 2015; Jentoft, 2007). Yet many clam fisheries have contracted markedly in recent decades, with closures and reduced availability reported for key target species and fishing grounds (Baeta et al., 2014, 2021).

This article examines Spanish clam fisheries as a case of cumulative stressors and persistent governance constraints. By focusing on how fishers, fishers’ guild leaders, and regional managers interpret ecological change and management effectiveness, the study complements stock- and landings-based assessments with systematic insight into mechanisms, institutional bottlenecks, and feasible pathways for reform (Davis & Wagner, 2003; Karnad, 2022).

### 1.1. Small-scale shellfisheries in nearshore social-ecological systems

SSF provide food, employment, and cultural value, but often operate with limited buffers against economic shocks and environmental extremes, particularly where livelihoods are locally dependent, and diversification options are constrained (FAO, 2024). Governance arrangements shape how rules are designed, who participates in decisions, and how costs and benefits are distributed — features that influence legitimacy and, ultimately, compliance. For clam fisheries, this governance setting is especially consequential. Habitat suitability depends on sediment processes and water quality, and productivity can be strongly affected by episodic disturbances. These characteristics increase sensitivity to cumulative pressures: chronic extraction can reduce resilience, while short-term shocks can then trigger abrupt declines or prolonged non-recovery (Baeta et al., 2014, 2021). In parallel, contested coastal space can intensify conflicts and complicate fisheries management, especially when fisheries are displaced or marginalised in spatial decisions (Bennett et al., 2015; Jentoft, 2007).

### 1.2. Spanish clam fisheries: species, fleets, and emerging pressures

Within Spanish SSF, clam fisheries have historically supported coastal employment and seasonal income along both the Mediterranean and southern Atlantic coasts (Andalusia). The principal target species include wedge clam (*Donax trunculus*), striped venus clam (*Chamelea gallina*), and smooth clam (*Callista chione*). These fisheries are strongly local in their spatial footprint, typically operating in shallow, nearshore habitats where environmental conditions can vary sharply over short distances and where human uses overlap.

Available evidence suggests that several Spanish clam fisheries have experienced long-term declines and heightened management difficulty (Baeta et al., 2026). For example, research in the NW Mediterranean documents the decline of *C. chione* associated with interacting pressures and reduced recovery potential (Baeta et al., 2014). Meanwhile, longer-term analyses highlight persistent vulnerability and management challenges in clam fisheries under changing environmental and governance conditions (Baeta et al., 2018, 2026). Additional work has documented the collapse and lack of recovery of the *D. trunculus* fishery in the Gulf of Valencia following its closure (Escrivá et al., 2021).

Stakeholder accounts in the present study point to a range of perceived drivers, including environmental variability and extremes, habitat alteration linked to coastal works and sediment dynamics, water quality pressures, episodic disease or mortality events, and fishing pressure amplified by illegal extraction and informal marketing. Understanding how these pressures combine—and how governance systems enable or constrain response—is central to designing policy that can reduce collapse risk and sustain livelihoods.

### 1.3. Local ecological knowledge for diagnosing cumulative stressors and governance constraints

Local ecological knowledge (LEK) has become increasingly recognised as a valuable source of information for understanding ecological change and informing fisheries management, particularly in data limited contexts and nearshore settings where conditions are spatially heterogeneous, and monitoring can be sparse (Davis & Wagner, 2003; Gerhardinger et al., 2009). When elicited through structured methods, LEK can help document ecological changes and provide insights into their potential drivers (Ainsworth, 2011; Cavole et al., 2020). Recent marine policy contributions also emphasise the value of integrating LEK to inform fisheries management and conservation strategies, particularly in data-limited contexts (Gomes et al., 2025).

In Spain, clam fisheries are governed within a decentralised administrative context, with management authority distributed across Autonomous Communities (AACC). Decentralised governance can enable context specific regulation, but it can also create fragmented rules, uneven enforcement capacity, and coordination gaps across adjacent fishing grounds. Such institutional “*limits of governability*” are particularly salient for small scale, nearshore fisheries where compliance depends on both credible enforcement and perceived fairness (Jentoft, 2007; Nielsen, 2003; Ostrom, 1990). LEK is therefore well-suited to diagnose not only ecological trajectories but also the institutional constraints that shape whether management measures are timely, legitimate, and enforceable.

### 1.4. Study aims and analytical expectations

In this context, we examine (i) how stakeholders interpret ecological change and the build-up of cumulative stressors, and (ii) which institutional conditions they see as shaping management outcomes in Spanish clam fisheries. We pursue three objectives: (1) to document perceived ecological trends and stressors affecting the main clam fisheries across regions and species; (2) to identify governance and management constraints—particularly around monitoring, enforcement, participation, and cross-scale coordination—that stakeholders associate with (in)effectiveness; and (3) to compare perspectives across regions and actor groups, highlighting where explanations and proposed solutions converge or diverge.

Guided by these objectives, we examine three analytical expectations using a comparative qualitative approach. H1 (Multiple perceived stressors): stakeholder accounts identify environmental, habitat, biological, and fishing/socio-economic pressures associated with clam decline. H2 (Governance constraints): stakeholder accounts identify perceived management limitations and institutional constraints, including fragmented authority, uneven implementation of rules, and limitations in monitoring and enforcement. H3 (Context dependence): stakeholder priorities concerning perceived drivers and management responses vary across study areas. These expectations guide the descriptive analysis; they are not treated as formal tests of causal relationships.

## 2. Materials and methods

We combined contextual analysis of the institutional setting with semi-structured interviews to examine how fishers, fishers’ guild leaders, and regional managers interpret (i) ecological change and multiple perceived stressors and (ii) the governance conditions shaping management outcomes in Spanish clam fisheries. The design follows common qualitative approaches in marine policy and SSF research, where LEK is used to illuminate mechanisms and institutional constraints in data-limited or highly heterogeneous coastal systems.

### 2.1. Institutional and governance context

The governance of clam fisheries in Spain was decentralised in the early 1980s, when the AACC assumed responsibility for fisheries management within their areas of competence, including fishing in internal waters and shellfishing. The AACC represent regional administrative jurisdictions that vary considerably in coastline length and number of fishing grounds (**Fig. 1**). As a result, each AACC has developed its own regulatory framework, leading to regional differences in licensing systems, permitted fishing gears, fishing areas and seasons, minimum landing sizes, catch and effort restrictions, and monitoring and enforcement. This institutional heterogeneity provides the broader context for analysing stakeholder perceptions of governance effectiveness, coordination, and legitimacy.

**Figure 1.**
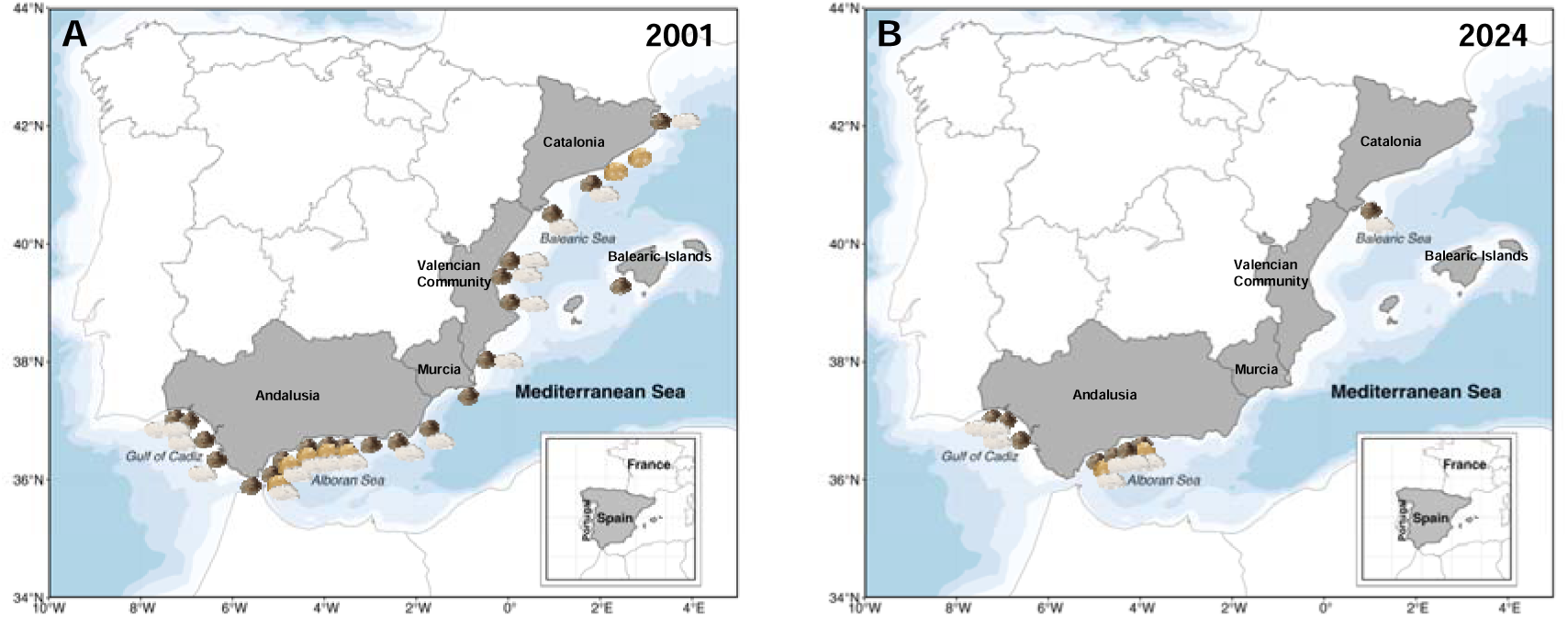
Geographic distribution of clam fisheries (*Chamelea gallina*, *Donax trunculus*, and *Callista chione*) along Spain’s Mediterranean and southern Atlantic coasts. **(A)** Data for 2001, compiled from the inventory (Alarcón, 2001), relevant legal documents for the Mediterranean coast, and landing notes for Atlantic Andalusia (Andalucía, 2025). **(B)** Data for 2024, based on landing notes from Catalonia and Andalusia (Andalucía, 2025; Generalitat de Catalunya, 2025). Each shell icon indicates active commercial fisheries targeting these species (*D. trunculus*, *C. gallina*, and *C. chione*) in each area. The sampled areas correspond to MED (Catalonia, Valencian Community, Balearic Islands, and Murcia), ALB (Mediterranean Andalusia), and ATL (Atlantic Andalusia). Overall, the maps show a pronounced contraction of fishing activity over the past two decades, with most fishing grounds closed or inactive by 2024.

The study encompasses five AACC (Catalonia, Valencian Community, Balearic Islands, Murcia, and Andalusia), represented by six sampled coastal strata because Andalusia was considered separately for its Mediterranean and Atlantic coasts. For comparative analyses, these strata were grouped into three broader coastal regions: MED (Catalonia, Valencian Community, Balearic Islands, and Murcia), ALB (Mediterranean Andalusia), and ATL (Atlantic Andalusia). The study focuses on AACC that have historically supported clam fisheries (**Fig. 1**). Galicia was excluded because the focal species—*D. trunculus*, *C. gallina*, and *C. chione*—have not represented major fisheries there; Galician bivalve production mainly targets other taxa (*e.g*., *Cerastoderma edule*, *Pecten maximus*, *Ruditapes decussatus*, and *Ruditapes philippinarum*).

### 2.2. Study area and biogeographical framing

To provide an environmental and biogeographical frame that better reflects clam population distributions than administrative boundaries, we aggregated interview and landings context into three coastal regions: (i) the NW Mediterranean (Catalonia, Valencian Community, Balearic Islands and Murcia), (ii) the Alboran Sea (Mediterranean coast of Andalusia) and (iii) the Gulf of Cádiz (Atlantic coast of Andalusia). This framing supports consistent cross regional comparison while recognising that nearshore ecological dynamics and pressures differ across oceanographic settings.

### 2.3. Fishery and temporal context

Long-term landings context was used to situate interview narratives and to support a coherent reading of perceived ecological trajectories (**Fig. 2**; data exclude Galicia). In the NW Mediterranean, landings of *C. gallina*, *D. trunculus*, and *C. chione* (**Fig. 2A-C**) began a progressive decline from the 1990s, culminating in successive closures of most fishing grounds. *C. gallina* fisheries in Murcia and the Balearic Islands were closed in 2002; fishing activity targeting *C. chione* was abandoned in Catalonia in 2007; and in 2015, all *D. trunculus* fisheries in the Valencian Community and most of those operating in Catalonia were closed due to low catches (Baeta et al., 2026).

**Figure 2.**
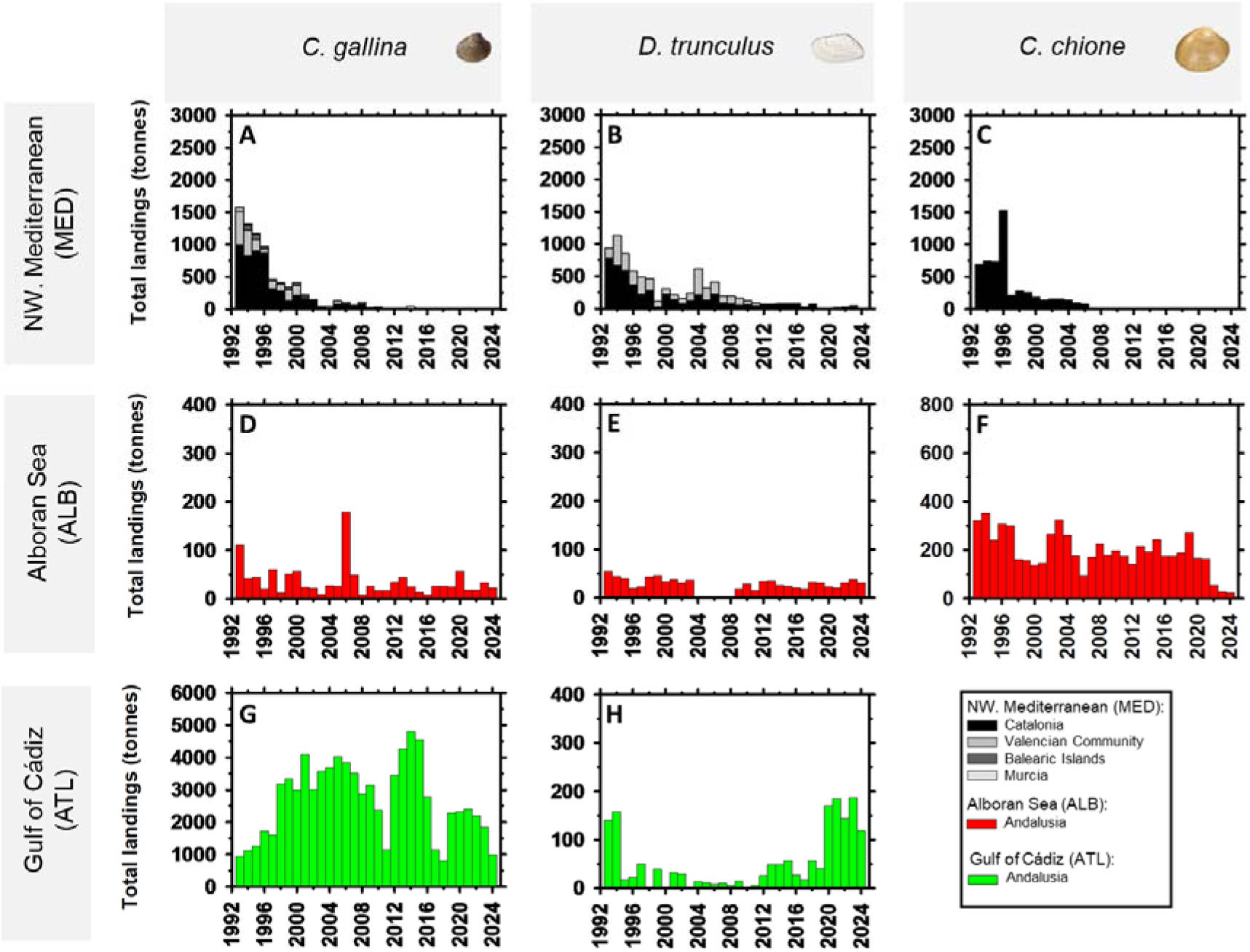
Temporal trajectories of official landings of the three main clam species across Spanish coastal regions (1993–2024). Total annual landings (tonnes) are shown for *Chamelea gallina* (**A, D and G**), *Donax trunculus* (**B, E and H**), and *Callista chione* (**C and F**) across three regions: NW Mediterranean (MED; **A–C**), Alboran Sea (ALB; **D–F**), and Gulf of Cádiz (ATL; **G–H**). Years without visible bars indicate zero reported landings rather than unavailable data. Within the NW Mediterranean, stacked grey bars distinguish landings from Catalonia, Valencian Community, Balearic Islands, and Murcia, while red and green bars represent Andalusian landings from the Alboran Sea and Gulf of Cádiz, respectively. Note that y-axis scales differ among panels.

In the Alboran Sea, landings of *C. gallina* and *D. trunculus* show marked interannual variability since the 1990s, characterised by alternating phases of decline and short-lived recoveries (**Fig. 2D-E**). In contrast, *C. chione* remained relatively stable for nearly three decades, with a notable drop only in the most recent years (**Fig. 2F**). The region has also experienced a gradual reduction in active fishing grounds, particularly along its northern sector.

In the Gulf of Cádiz, on the Atlantic coast, *C. gallina* supports the most productive clam fishery in Spain, with high but fluctuating landings and alternating periods of sharp declines and strong recoveries since the 1990s (**Fig. 2G**). *D. trunculus* also shows pronounced interannual variability, with an apparent long-term decline followed by a partial increase in reported landings from the 2010s onwards. However, changes in data recording may have contributed to this apparent increase, although the available landing data do not allow this possibility to be distinguished from a true biological change (**Fig. 2H**). Unlike the Mediterranean region, *C. chione* is not commercially harvested in Gulf of Cádiz.

### 2.4. Interview design

We used semi-structured interviews to elicit stakeholder perspectives on two core themes: (1) perceived drivers of decline and ecological change, and (2) clam fishery governance and management practice (see **Supplementary Material** I for the complete interview protocol). The protocol combined structured questions used to characterise participants and the different types of clam fisheries in each AACC with predominantly open-ended questions addressing fishery trajectories, perceived drivers of decline, fishing effort, management, governance, monitoring and enforcement, informal practices, and conflicts. No Likert-type or other ordinal response scales were used, although some questions requested quantitative or semi-quantitative information. The semi-structured format was chosen to balance open discussion with consistent thematic coverage across AACC and actor groups, enabling comparison while allowing participants to raise locally salient stressors and governance concerns.

### 2.5. Sampling and participants

Between April 2024 and August 2025, we conducted 94 interviews across coastal AACC with historical clam fisheries activity in the Spanish Mediterranean and Atlantic (excluding Galicia). Participants were recruited through purposive sampling, complemented by snowball referrals, to include key stakeholder categories and fishing modalities. Interviewees included active and former fishers, fishers’ guild (*cofradía*) leaders, and regional fisheries managers. Eligible participants were required to have direct experience with, or professional knowledge of, clam fisheries in the study area. Initial contacts were established through fishers’ guilds and associations, which facilitated contact with active and former fishers; additional participants were identified through referrals from interviewees. Recruitment continued throughout the study period and ceased when no further accessible eligible participants could be recruited.

The geographical distribution of interviews was purposive, with greater interview coverage sought in areas with historically more important clam fisheries, particularly Catalonia, the Valencian Community, and Andalusia. The sample was not designed to be proportional to the number of fishers, vessels, fishing effort, or recorded landings in each AACC.

### 2.6. Data collection procedures

Interviews typically lasted 30-60 minutes and were conducted in Spanish. All interviews followed the same core semi-structured interview framework and were conducted by the same interviewer (MAS) to maintain consistency in the interview procedure across study areas. Most interviews were conducted remotely, via video call, to accommodate participants’ availability and geographic dispersion. Two interviews in Catalonia were conducted face-to-face at Vilanova Port (Barcelona). The interviewer was previously acquainted with four participants interviewed in Catalonia, but not with participants from the other study areas. Detailed notes were taken during the interviews, and all interviews were audio-recorded with participants’ consent and subsequently transcribed. Transcripts were first reviewed against the audio recordings by MAS. Transcripts were analysed in Spanish, and only quotations selected for inclusion in the manuscript were translated into English by MB.

Interview prompts covered: perceived causes of declining landings; access and appropriation rules; management measures and their timing; participation in decision-making; monitoring and enforcement capacity; informal practices (*e.g*., off-auction sales and poaching); and conflicts within the sector and with other coastal users. Illustrative quotations were selected from the coded material when they expressed a recurrent theme particularly clearly. Quotations used in the Results are attributed only by region and role (fisher, guild leader, or regional manager) to preserve anonymity.

### 2.7. Ethics and confidentiality

All participants provided informed consent before participation. Participation was voluntary, and interviewees could decline to answer questions or withdraw at any point. To protect confidentiality, personal identifiers were removed from notes and transcripts, and quotes are reported using broad descriptors (region and stakeholder profile) rather than names, ports, or vessels. Recordings and transcripts were stored separately, and access was restricted to the researchers responsible for their handling. No formal approval from an institutional research ethics committee was sought for this study.

### 2.8. Qualitative analysis

We analysed notes and transcripts using thematic coding that combined deductive categories derived from the interview guide with inductive codes emerging from participants’ narratives. Coding focused on: (i) ecological and environmental stressors; (ii) fishing and market dynamics; (iii) governance arrangements, rules-in-use and participation; (iv) monitoring, enforcement and informal practices; and (v) conflict and cross-sector interactions in nearshore space. All transcripts were coded by the first author (MB). The coding unit was a text segment containing information relevant to a given theme. Initial codes combined themes derived from the interview guide with additional codes identified during transcript review; related codes were subsequently consolidated into higher-order thematic categories. For frequency summaries, repeated references to the same code by the same respondent were counted once, whereas a respondent could contribute to more than one code when multiple themes were mentioned. For the retrospective estimates of catch distribution presented in **Table 2**, respondents estimated the percentages of total catches attributable to poaching and to off-auction sales by licensed fishers. The percentage marketed through official fish auctions was calculated as the remainder to 100%. Passages whose meaning was insufficiently clear to support assignment to a specific code were left uncoded. Missing responses were distinguished from explicit statements of uncertainty or lack of knowledge, which were classified as “no opinion” where applicable. H1 was evaluated descriptively through the thematic coding of perceived drivers of decline, their grouping into broader stressor categories, and the frequency with which respondents mentioned each coded sub-driver across study regions (**Table S2, Supplementary Material II**). For summaries of perceived drivers, counts were based on valid responses within each AACC (Catalonia, n = 22; Valencian Community, n = 22; Balearic Islands, n = 4; Murcia, n = 4; Mediterranean Andalusia, n = 15; Atlantic Andalusia, n = 21; total valid responses, n = 88). Missing responses (n = 6) were excluded from these denominators, whereas explicit “no opinion” responses were retained as valid responses. The analysis was intended to identify patterns in stakeholder perceptions rather than to statistically test causal relationships among stressors. H2 was evaluated descriptively by examining stakeholder assessments of management effectiveness together with recurrent themes concerning governance arrangements, monitoring, enforcement, participation, and the implementation of rules. These themes are interpreted descriptively and are not used to infer causal relationships between institutional conditions and management performance.

Open-ended responses on management and governance were coded according to the assessments expressed by respondents into the 8 management and 7 governance categories shown in **Fig. 4**. These categories were subsequently grouped into three broader classes: positive, negative/insufficient, or no opinion. “Partially effective” was classified as positive, because it acknowledged some degree of effectiveness, whereas “improved, not enough” and “partially effective, but insufficient” were classified as negative/insufficient because respondents explicitly considered the improvements or outcomes inadequate. The same classification criteria were applied to management and governance assessments across all study areas. Finally, H3 was evaluated descriptively by comparing the salience and content of coded themes across AACC while the distribution of fishing modalities and actor roles across study areas was used to contextualise these comparisons (**Table S1, Supplementary Material II**). These comparisons were used to identify contextual variation in stakeholder perceptions rather than to test causal effects of nearshore dynamics or other environmental and fisheries characteristics.

Qualitative data were organised and coded in ATLAS.ti (version 25) (ATLAS.ti Scientific Software Development GmbH, 2025). To support transparent synthesis, we generated descriptive frequency summaries of recurrent themes (used to indicate salience, not statistical inference) and grouped codes into higher-order categories corresponding to ecological, governance, and socio-economic dimensions. Visual summaries were produced using SankeyMatic and SigmaPlot 16 (descriptive plots) (Systat Software, 2022).

### 2.9. Rigour and reporting

To enhance rigour, we (i) used a shared interview guide to ensure thematic comparability across regions, (ii) documented analytic decisions through iterative codebook refinement, and (iii) triangulated stakeholder accounts against the landings and institutional context presented in **Figs. 1**–**2** and associated text. We report the methods in sufficient detail to support assessment of transferability to other nearshore bivalve fisheries facing multiple stressors and multi-level governance challenges.

## 3. Results

Results are organised around (i) the profile of interviewees, (ii) perceived ecological trajectories and drivers of decline, and (iii) institutions and rules-in-use shaping management outcomes, including informal practices, participation, enforcement, and conflicts in nearshore space.

### 3.1. Interviewee profile and coverage

The 94 interviewees included stakeholders directly involved in clam fisheries across the Spanish Mediterranean and the Andalusian Atlantic coast (**Table 1**). Respondents were predominantly male (97.87%); two participants were women (2.13%). Ages ranged from 31 to 88 years (mean = 53), and interviewees reported long professional trajectories, typically spanning three decades.

**Table 1.** Socio-demographic and professional characteristics of interviewees (n = 94), including age, gender, Autonomous Community (AACC) and zone, role in the fishery (vessel-based mechanised clam dredges; on-foot hand-operated dredges; vessel-towed dredges 1, “*rastrell de cadenes*”; vessel-towed dredges 2, “*rastros*”; vessel-based hydraulic dredges; fishers’ guild leaders; and regional managers), occupational status, and years of fishing experience. Values are reported as counts and percentages. *Occupational status and years of fishing experience were calculated for participants classified as fishers (n = 70); fishers’ guild leaders and regional managers were excluded from these variables.

| Variable | Category | N (94) | Frequency (%) |
| --- | --- | --- | --- |
| Age | ≤ 40 | 3 | 3.19 |
|  | 41-50 | 9 | 9.57 |
|  | 51-60 | 52 | 55.32 |
|  | ≥ 61 | 30 | 31.91 |
| Gender | Male | 92 | 97.87 |
|  | Female | 2 | 2.13 |
| AACC / Zone | Catalonia | 24 | 25.53 |
|  | Valencian Community | 22 | 23.40 |
|  | Balearic Islands | 4 | 4.26 |
|  | Murcia | 4 | 4.26 |
|  | Andalusia Med. | 16 | 17.02 |
|  | Andalusia Atl. | 24 | 25.53 |
| Role | Mechanised clam dredges fishers | 36 | 38.30 |
|  | Hand-operated dredges fishers | 17 | 18.09 |
|  | Towed dredges 1 fishers | 5 | 5.32 |
|  | Towed dredges 2 fishers | 2 | 2.13 |
|  | Hydraulic clam dredges fishers | 10 | 10.64 |
|  | Fishers’ guild leaders | 19 | 20.21 |
|  | Regional managers | 5 | 5.32 |
| Current occupational status of fishers (N=70) | Active fishers | 34* | 48.57 |
|  | Other activity | 23* | 32.86 |
|  | Retired | 13* | 18.57 |
| Years of fishing experience (N=70) | ≤ 20 | 9* | 12.86 |
|  | 21-30 | 14* | 20.00 |
|  | 31-40 | 28* | 40.00 |
|  | ≥ 41 | 19* | 27.14 |

Nearly half of the fishers were active (48.57%), while 32.86% had transitioned to other occupations and 18.57% were retired, providing both contemporary and retrospective accounts of fishery change. In terms of roles, mechanised dredge fishers formed the largest group (38.30%), followed by fishers’ guild (*cofradía*) leaders (20.21%), hand-operated dredge fishers (18.09%), hydraulic dredge fishers (10.64%), regional managers (5.32%), vessel-towed dredges 1 fishers (5.32%), and vessel-towed dredges 2 fishers (2.13%).

Interviews were distributed across all Mediterranean AACC with historical clam fisheries and both Andalusian coasts. Catalonia (n = 24), Atlantic Andalusia (n = 24) and the Valencian Community (n = 22) accounted for most interviews, with additional coverage of Mediterranean Andalusia (n = 16) and smaller samples from the Balearic Islands (n = 4) and Murcia (n = 4). These two small regional samples provide limited-coverage accounts rather than representative regional estimates. The distribution of participants by study area and actor role/fishing modality is provided in **Table S1** (**Supplementary Material II**). Broader socioeconomic and cultural characteristics of participants were not systematically collected and therefore were not used to stratify the sample.

### 3.2. Perceived trajectories of decline

Across regions and actor groups, interviewees described long-term contraction of clam fisheries, typically as a gradual process rather than a single collapse event. Overall, 66 respondents (70.2%) characterised decline as progressive, while 28 (29.8%) reported abrupt collapses in particular grounds or periods. Narratives frequently linked these trajectories to weak or absent recovery following shocks, with closures and loss of productive beds described as cumulative over time.

### 3.3. Perceived drivers of decline and multiple stressors

Stakeholder accounts converged on a multifactorial explanation of decline, grouped into five broad categories: (i) climate-related stressors, (ii) anthropogenic alteration of coastal habitats, (iii) biological stressors, (iv) governance and institutional shortcomings, and (v) fishing pressure coupled with socio-economic drivers (**Fig. 3**; **Table S2**). Respondents described these pressures through specific observations and explanations rather than general statements. Climate-related pressures included high temperatures, drought or reduced rainfall, storms, and general climate change. Coastal alteration was described through sediment-supply changes, beach nourishment and dredging, wastewater and agricultural runoff, and pressures associated with mass tourism. Biological stressors included pathologies and diseases. Fishing and socio-economic pressures included overfishing and poaching.

**Figure 3.**
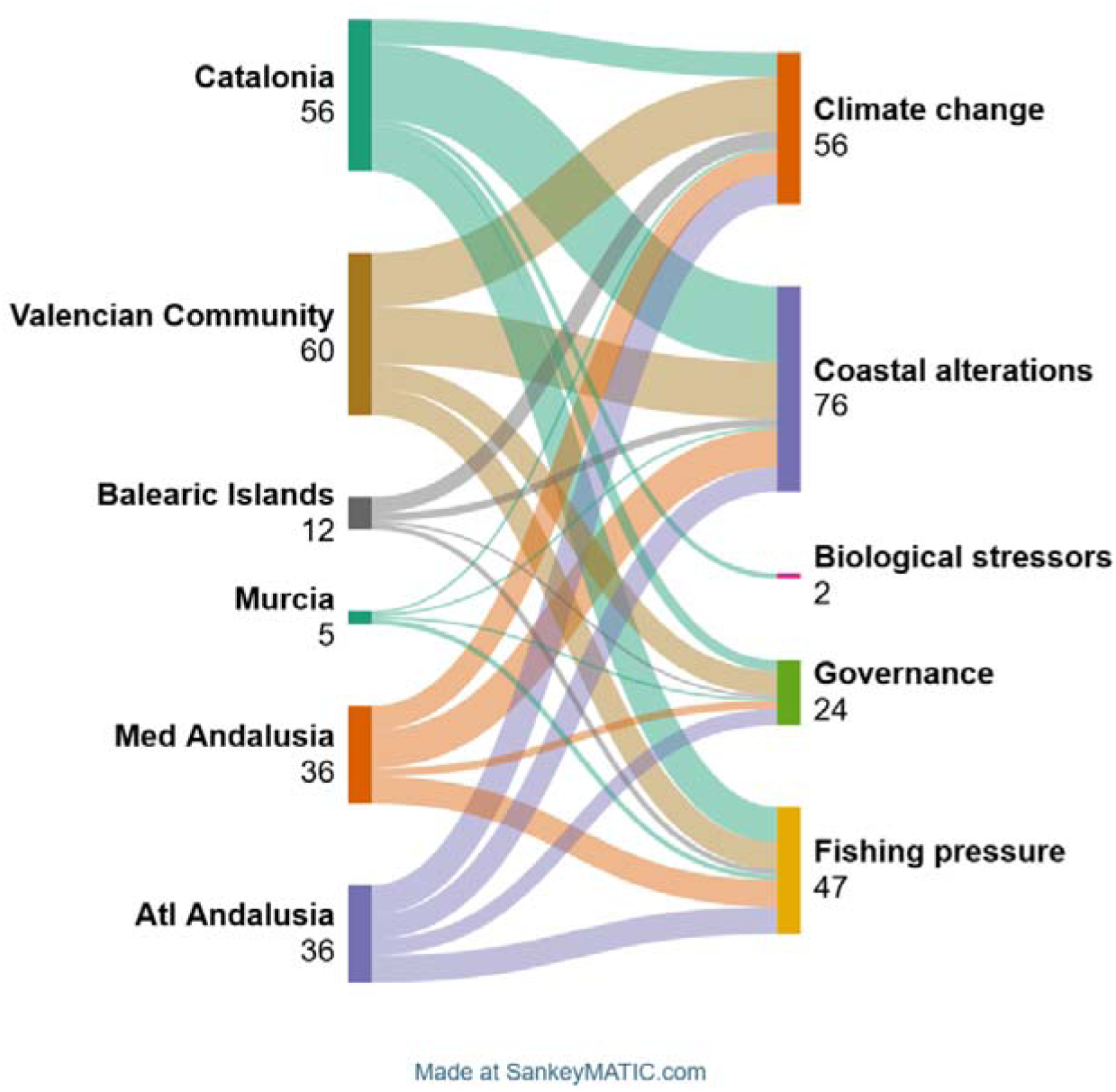
Sankey diagram showing stakeholder perceptions of the main drivers of clam-fishery decline across the study areas (205 sub-diver mentions from 88 valid responses among 94 interviews). Flows connect each study area (left) to a driver category (right): climate change, coastal alterations, biological stressors, governance and institutional factors, and fishing pressure. Flow widths represent raw counts of respondent-level sub-driver mentions aggregated within each driver category and are not normalised by the number of valid responses in each study area. Repeated mentions of the same sub-driver by the same respondent were counted once, whereas a respondent could contribute more than once to a broader driver category when different sub-drivers within that category were mentioned. The diagram describes the distribution of themes in the interview material; it does not establish causal interactions among stressors. Sub-driver mention counts by study area are reported in **Table S2 (Supplementary Material II).**

Perceived driver salience varied across study areas, with the relative frequency of individual sub-drivers differing among AACC, with the number of mentions of individual sub-drivers differing among areas (**Table S2, Supplementary Material II**). These descriptive contrasts should be interpreted in the light of differences in interview coverage and participant composition across study areas. These patterns are visualised in the Sankey diagram (**Fig. 3**), where flow widths represent raw counts of respondent-level sub-driver mentions aggregated within each driver category rather than within-area frequencies.

### 3.4. Perceived effectiveness of management and governance

Most interviewees characterised clam fishery management as ineffective or insufficient (n = 60; **Fig. 4A**). Respondents commonly reported that management had either never been effective (n = 25) or had been effective only in the past under stronger local self-management arrangements but not after responsibility shifted to regional governments (n = 20). A smaller number suggested that management has deteriorated over time (n = 7), while 8 respondents reported that management has improved but remained insufficient. In contrast, 29 interviewees described management as effective to varying degrees, including consistently effective (n = 10), partially effective (n = 8), or previously ineffective but now effective (n = 11). Three respondents reported no opinion. Regional management assessments also varied across study areas (**Fig. 4B**). Three of four interviewees from the Balearic Islands reported positive assessments, whereas negative or insufficient assessments predominated in the Valencian Community, Murcia, and both Andalusian coasts. Catalonia showed a more balanced distribution of positive and negative assessments.

**Figure 4.**
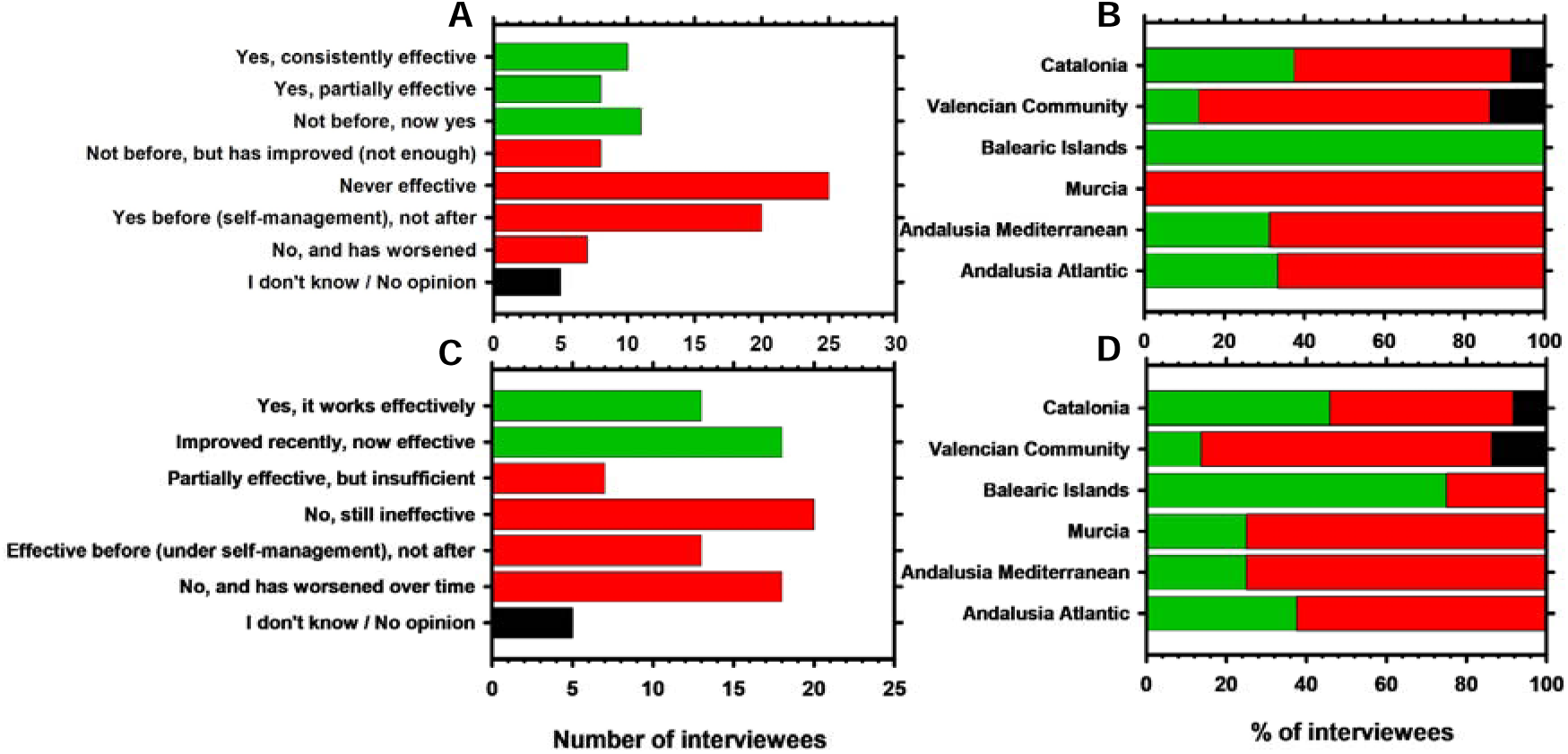
Stakeholder perceptions of clam fishery management and governance effectiveness (n = 94 interviewees). **(A)** Overall perceptions of management effectiveness: green bars indicate positive assessments, red bars indicate negative or insufficient assessments, and the black bar indicates no opinion. **(B)** Management effectiveness across the six study areas: bars show the percentage of interviewees within each area reporting positive assessments (green), negative or insufficient assessments (red), or no opinion (black). **(C)** Overall perceptions of governance effectiveness, using the same classification as in panel A. **(D)** Governance effectiveness across the six study areas, expressed as the percentage of interviewees within each study area using the same classification as in panel **B**.

Governance assessments showed a broadly similar overall pattern (**Fig. 4C**), with 58 interviewees expressing critical or insufficient assessments and 31 describing governance as effective or recently improved; 5 reported no opinion. Regional governance assessments also varied across study areas (**Fig. 4D**). Three of the four interviewees from the Balearic Islands reported positive assessments, whereas negative or insufficient assessments predominated in the Valencian Community, Murcia, and both Andalusian coasts. In Catalonia, positive and negative /insufficient assessments were equally frequent (n =11).

Across interviews, explanations for perceived management and governance shortcomings most frequently referred to lack of coordination among actors, limited influence of fishers in decision-making, and implementation gaps between formal rules and practice.

### 3.5. Access, informal practices, and traceability

Interviewees emphasised that formal licensing and modality-based access rules often diverged from practices on the ground. Two informal practices were repeatedly highlighted as shaping both market dynamics and perceptions of legitimacy: off-auction sales and poaching.

Many respondents acknowledged that off-auction sales have occurred historically and, while perceived to have declined, were still reported in some areas. Interviewees described economic incentives (“*they pay more*”) and the imposition of purification requirements as key drivers of informal marketing, alongside attempts to commercialise undersized clams outside official control points. In contrast, some respondents reported that off-auction sales had not occurred in their areas.

Poaching was frequently referenced by interviewees and described as a longstanding issue, particularly in the wedge clam fishery (*D. trunculus*) associated with hand-operated dredges and beach-based activity. Respondents distinguished between professional illegal extraction and diffuse harvesting by beachgoers, especially during summer. Several stressed that enforcement is difficult because some offenders have limited resources, and penalties may be ineffective where insolvency is declared.

Perceptions of informal activity also varied by species and region (**Table 2**). For *D. trunculus*, interviewees described high levels of off-auction sales and poaching in the 1980s-1990s, with strong declines in Catalonia and Mediterranean Andalusia during the 2010s, alongside an increasing share attributed to auction sales. In contrast, informal channels were reported to persist in the Valencian Community and Atlantic Andalusia, including interviewee estimates indicating substantial illegal activity in areas where the fishery was officially closed. For *C. gallina*, interviewees generally characterised first sale through auctions as the dominant pathway since the 2000s across most regions, with only small fractions sold informally; poaching was described as limited and primarily historical in Andalusia.

**Table 2.** Perceived distribution of clam landings based on retrospective estimates provided by interviewees. Values represent the mean of individual estimates for each region and decade and are shown as % auction / % off-auction / % poaching. The number of valid interviewees contributing to each estimate is given in parentheses. Auction sales represent the remaining proportion after accounting for the estimated proportions attributed to off-auction sales and poaching, so that each triplet sums to 100%. Mean percentage estimates were rounded to the nearest whole number. Dashes (--) indicate that no fishery/resource was reported for that species, region and decade, whereas zero values indicate an estimated absence of the corresponding component. Estimates may nevertheless be reported for officially closed fisheries where interviewees reported continued illegal harvesting (poaching). These values represent stakeholder perceptions rather than official statistics. **(A)** *D. trunculus*; this species has not been fished in either Murcia or the Balearic Islands. **(B)** *C. gallina*.

| (A) Region / Decade | 1980s | 1990s | 2000s | 2010s | 2020s |
| --- | --- | --- | --- | --- | --- |
| Catalonia | 20 / 50 / 30 (6) | 40 / 50 / 10 (7) | 60 / 30 / 10 (8) | 75 / 20 / 5 (8) | 90 / 5 / 5 (6) |
| Valencian Community | 20 / 50 / 30 (6) | 40 / 50 / 10 (6) | 70 / 20 / 10 (9) | 70 / 25 / 5 (9) | 0 / 0 / 100 (5) |
| Andalusia Med. | 20 / 50 / 30 (5) | 50 / 40 / 10 (5) | 60 / 30 / 10 (7) | 85 / 10 / 5 (7) | 90 / 5 / 5 (7) |
| Andalusia Atl. | 20 / 50 / 30 (8) | 40 / 30 / 30 (8) | 50 / 30 / 20 (9) | 60 / 20 / 20 (9) | 85 / 5 / 10 (9) |

| (B) Region / Decade | 1980s | 1990s | 2000s | 2010s | 2020s |
| --- | --- | --- | --- | --- | --- |
| Catalonia | 75 / 25 / 0 (5) | 70 / 30 / 0 (6) | 70 / 30 / 0 (6) | 95 / 5 / 0 (8) | 5 / 5 / 0 (6) |
| Valencian Community | 75 / 25 / 0 (4) | 70 / 30 / 0 (6) | 80 / 20 / 0 (9) | 90 / 10 / 0 (9) | -- |
| Balearic Islands | -- | 70 / 30 / 0 (3) | 70 / 30 / 0 (3) | -- | -- |
| Murcia | 80 / 20 / 0 (3) | 90 / 10 / 0 (3) | 100 / 0 / 0 (3) | -- | -- |
| Andalusia Med. | 60 / 35 / 5 (6) | 70 / 30 / 0 (5) | 85 / 15 / 0 (7) | 90 / 10 / 0 (8) | 5 / 5 / 0 (5) |
| Andalusia Atl. | 60 / 35 / 5 (8) | 70 / 30 / 0 (8) | 90 / 10 / 0 (6) | 95 / 5 / 0 (8) | 90 / 10 / 0 (7) |

**Note:** For the Balearic Islands and Murcia, estimates for the 2000s cover a decade during which the respective fisheries were closed in 2002; these retrospective estimates should not be interpreted as evidence of authorised fishing throughout that decade.

### 3.6. Governance frameworks and rules-in-use

Stakeholders described a governance trajectory from predominantly bottom-up arrangements, centred on guild self-regulation and LEK, towards more formalised, top-down management following the transfer of competences to AACC in the 1980s. Respondents reported that minimum landing sizes, seasonal closures, and gear restrictions became codified through decrees during the 1990s, while more restrictive measures (*e.g*., total allowable catches, quotas, and first-sale requirements) were increasingly introduced from the 2000s onwards, which interviewees often associated with visible declines. Since the 2010s, interviewees noted the creation of management plans and follow-up committees that reintroduced participatory elements in some regions, although decision-making was still widely perceived as centralised.

Across regions, clam fisheries management relied on a relatively consistent set of appropriation rules. The most frequently mentioned were fishing schedules (70 mentions), total allowable catches (66), technical restrictions on dredges and vessels (56), defined fishing zones (50), first-sale points (38), minimum landing sizes (19) and temporary closures (11). Interviewees also emphasised the continued importance of internal guild regulations that were often stricter than administrative rules, including screening at sales points for minimum size compliance (37 mentions), daily quotas (26), restrictions on fishing hours/days (26), rotation schemes (5) and limits on haul duration (3).

When asked which measures were effective in practice, respondents most frequently identified total allowable catches (31 mentions), technical restrictions on dredges and vessels (22 mentions), fishing schedules (21 mentions), and defined fishing zones (16 mentions) (**Fig. 5**). Among these technical restrictions, interviewees particularly highlighted engine-power regulations (7 mentions) and dredge mesh-size regulations (5 mentions). More generally, interviewees stressed that many rules were perceived as technically reasonable but undermined by weak enforcement, limited participation, and insufficient transparency.

**Figure 5.**
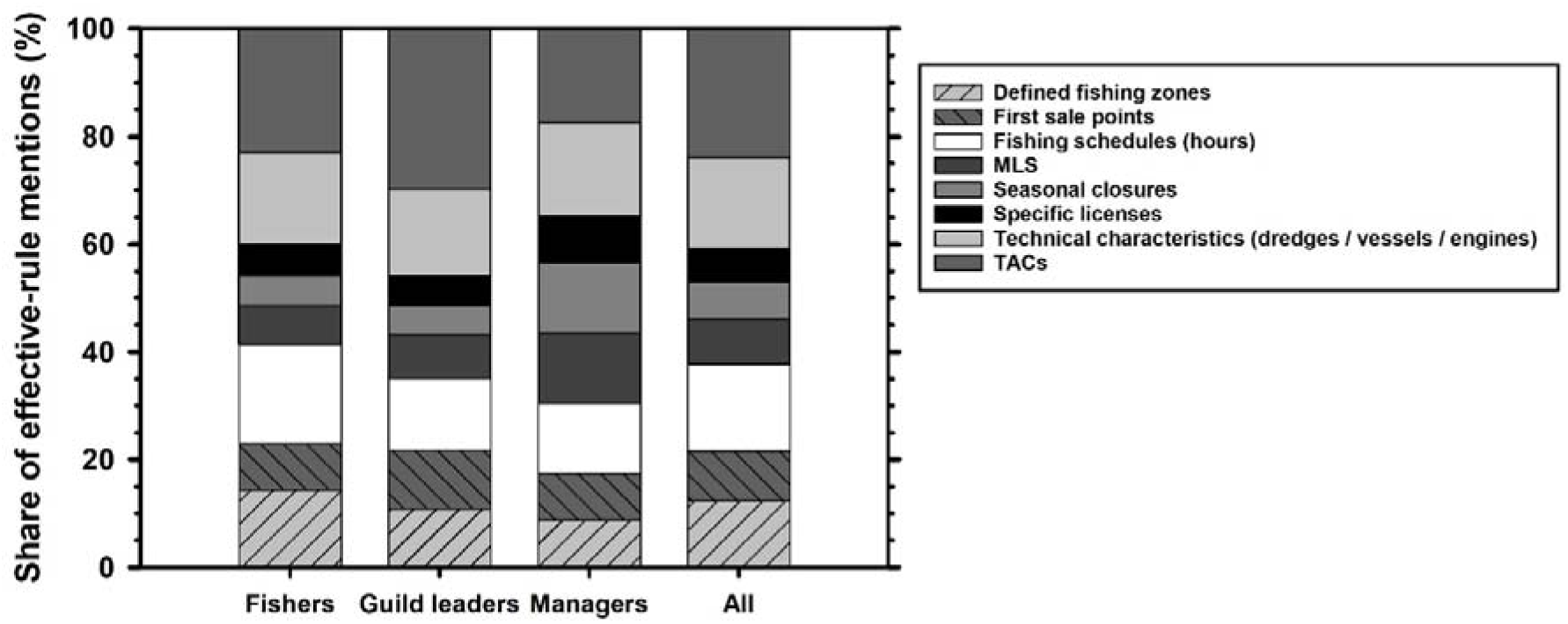
Stakeholder assessments of the effectiveness of appropriation rules, by actor group: fishers, fishers’ guild leaders, regional managers, and all respondents combined. Percentages represent the share of effective-rule mentions within each actor group, calculated as the number of mentions of each rule divided by the total number of effective-rule mentions in that group. The analysis included 46 fishers (70 mentions), 9 guild leaders (37 mentions), and 4 regional managers (23 mentions). Overall, the pooled sample comprised 59 respondents and 130 effective-rule mentions. The remaining 35 interviewees provided no information on the effectiveness of individual appropriation rules and were therefore excluded from this analysis. Repeated mentions of the same rule by the same respondent were counted once, whereas respondents could contribute to more than one rule when multiple effective measures were identified. The “All” category was calculated from pooled mentions across actor groups. MLS, minimum landing size; TACs, total allowable catches.

### 3.7. Participation, monitoring, and enforcement

Most interviewees reported that clam fishers remain organised primarily through local fishers’ guilds (*cofradías*), often with representation structured by fishing modality. Two notable exceptions were described: a cooperative established by mechanised dredge fishers on the Maresme coast (Catalonia) during the 1980s-2000s, and associations representing hand-operated dredge fishers on the Atlantic Andalusian coast. In some areas, interviewees also described supra-guild coordination arrangements spanning multiple guilds and modalities within shared fishing grounds.

Decision-making forums were reported to have evolved over time. Before the 1980s, meetings reportedly involved the Maritime Authority and the local guild leader. Following decentralisation, regional administrations assumed this role, meeting primarily with guild leaders, and later also with fisher representatives; scientific experts were increasingly involved from the 2000s onward. Interviewees described these meetings as structured and frequent where fisheries remain active, but they also stressed that not all participants felt heard, which they associated with frustration and limited influence over decision making.

Perceptions of ecological monitoring varied by AACC. Respondents described systematic monitoring as limited or discontinuous in Catalonia and the Valencian Community, often initiated after declines and later discontinued due to resource constraints. By contrast, interviewees in Murcia, the Balearic Islands, and both Andalusian coasts more frequently reported ongoing monitoring programmes. Across all regions, sanitary sampling to meet food safety requirements was described as routine, even where ecological monitoring was limited.

Perceived compliance with official regulations and internal guild rules was generally described as high among licensed vessel-based fishers, with infractions considered relatively uncommon. Sanctions—often associated with undersized catches or off-auction sales—were described as progressive and substantial. Respondents emphasised strong modality differences in enforceability: vessel-based fishers typically land through port auctions, which facilitates inspection, whereas on-foot fishers may transport catch by car, increasing opportunities for evasion. Enforcement responsibilities were attributed to multiple authorities (regional inspectors, national administration, SEPRONA, and occasionally guilds), and interviewees noted that differential enforcement across modalities can reinforce perceptions of unequal treatment.

### 3.8. Conflicts in nearshore space and conflict resolution

Interviewees reported diverse conflicts affecting clam fisheries, reflecting the crowded and multi-use nature of nearshore environments. The most frequently reported conflict was with beachgoers (34 mentions), especially during summer when recreational use overlaps with fishing activity. This was described as particularly acute for fisheries targeting *D. trunculus* in shallow areas that coincide with bathing beaches, where respondents perceived administrations as prioritising tourism through reduced working hours or restricted access. Some respondents also described diffuse informal harvesting by beach users as a form of small-scale poaching that they perceived could result in substantial cumulative removals.

Conflicts with regional administrations were also common (28 mentions), centring on the perceived burden of regulation and sanctions, the establishment of marine reserves, and mandatory depuration. Additional conflicts included coastal works and dredging for beach replenishment or harbour maintenance (11 mentions), professional poachers (6), recreational boating (5), and, less frequently, tensions with scientists advising administrations (4). Notably, 26 respondents reported no major conflicts affecting their activity.

In terms of resolution, respondents most often described direct dialogue as the first step (18 mentions), followed by mediation through regional administrations (15), local guilds (6), municipal councils (3), or the maritime authority (3). Nevertheless, many stakeholders emphasised that conflicts are difficult to resolve and may persist over time (22 mentions).

Conflicts among fishers were widely described as infrequent (91 respondents). Among the 88 respondents who provided information on how conflicts between fishers are typically resolved, regardless of their personal experience of such conflicts, 63 mentioned direct communication and 25 mentioned mediation by the fishing guild. Where disputes occurred, they were associated with competition for grounds, breaches of working hours, and illegal marketing. Some interviewees highlighted tensions between gear types exploiting the same species, and on-foot fisheries were occasionally singled out as being perceived as more prone to rule-breaking. In Atlantic Andalusia, several respondents reported that conflicts in the *C. gallina* fishery have increased in recent years, linked to stricter regulation, declining catches, and perceived unfair competition from imported products originating from fisheries subject to lower minimum landing sizes (MLS) than those applied in Spain. EU legislation established a minimum landing size (MLS) of 25 mm for *C. gallina* (Union, 2006). However, this threshold was reduced to 22 mm in specified Italian territorial waters through EU-approved derogations initially under the Commission Delegated Regulation (EU) 2016/2376 (European Commission, 2016) and subsequently maintained during the study period under Commission Delegated Regulation (EU) 2022/2587 (European Commission, 2022). In addition, in Turkey, a minimum landing size (MLS) of 17 mm has been reported for *C. gallina* (Ceylan, 2022; Scarcella & Mosteiro Cabanelas, 2016). These regulatory differences provide context for the market asymmetries interviewees perceived and the tensions reported by fishers operating under stricter national frameworks.

### 3.9. Integrated patterns across stressors and governance constraints

Across study areas, interviewees identified both perceived drivers of clam-fishery decline and difficulties involved in managing these fisheries. Reported themes included: environmental pressures, coastal alterations, biological stressors, and fishing-related pressures, together with limitations in monitoring, enforcement, and participation, as well as differences between formal rules and implementation in practice. However, these patterns were not uniform across the study areas. The relative importance of perceived stressors varied among study areas (**Table S2, Supplementary Material II**), while the distribution of fishing modalities and actor roles also differed among areas (**Table S1, Supplementary Material II**). Taken together, these accounts provide a descriptive overview of the perceived ecological pressures and management constraints reported across the clam fisheries examined in Spain. These descriptive patterns do not establish causal relationships or disentangle regional differences from variation in sample composition.

## 4. Discussion

This study used local ecological knowledge (LEK) to examine how fishers, fishers’ guild leaders, and regional managers interpret the decline of Spanish clam fisheries and the governance conditions shaping management outcomes. Across regions and actor groups, respondents identified multiple perceived pressures associated with clam declines, including environmental, habitat, biological, and fishing-related stressors, while also identifying governance constraints—fragmented authority, uneven rules among gears and territories, limited participation, and weak monitoring and enforcement—as recurrent limitations to management.

Two implications follow from these accounts. First, stakeholders identified climate variability, coastal development, and extraction as multiple perceived pressures, motivating a combined examination of ecological and institutional concerns (Baeta et al., 2014, 2018, 2021). Second, knowledge of ecological risks needs to be considered alongside institutional capacity, coordination, and legitimacy (Bennett et al., 2015; Jentoft, 2007; Ostrom, 1990). The following subsections distinguish stakeholder observations from interpretations in the literature and discuss options for more coherent coastal management.

### 4.1. Perceived pressures and possible mechanisms of decline

Most interviewees described gradual declines, while 28 of 94 reported abrupt declines in particular grounds or periods. These accounts can be considered alongside research on the long-term effects of dredging on bivalve stocks (Piersma et al., 2001). They are also consistent with the broader interpretation that chronic pressures may reduce recovery capacity (Baeta et al., 2014, 2018, 2021). However, the present interviews do not establish ecological thresholds or causal interactions.

Stakeholder narratives highlighted three broad groups of perceived pressures. First, chronic fishing pressure was frequently perceived as a background driver of decline, with interviewees referring to increasing fishing pressure and concentration of effort on remaining productive clam beds. These processes align with analyses of Spanish clam fisheries linking sustained fishing pressure to long-term declines in clam landings and stock condition (Baeta et al., 2014, 2018, 2021). In sedentary or low-mobility species, these patterns suggest that local depletion may be masked in aggregated statistics as effort redistributes spatially, delaying recognition of decline and encouraging reactive rather than preventive management.

Second, interviewees identified environmental variability and climate-related pressures, particularly high temperatures, drought or reduced rainfall, and storms, as perceived contributors to clam-fishery decline. This interpretation is consistent with literature showing that bivalve biomass and physiological condition can respond to variation in temperature and salinity, with responses depending on the timing and magnitude of environmental variation and on species-specific physiological sensitivity (Bento de Almeida et al., 2021; Soms-Molina et al., 2024). These perceived pressures may interact, but such interactions were not tested here. The consequences of fishing pressure may depend on whether environmental extremes coincide with spawning, settlement, or recovery periods; this remains a mechanism to investigate rather than a relationship established by the interviews.

Third, habitat alteration and water-quality degradation were repeatedly identified as perceived contributors to clam-fishery decline. Nearshore clams depend on sediment characteristics (mainly granulometry) and beach-shoreface processes; respondents identified sand dredging, beach nourishment, and other coastal alterations as perceived pressures on clam fisheries. Previous work has shown that such habitat alteration can reduce available clam habitat and concentrate fishing activity in the remaining grounds, thereby increasing fishing pressure (Baeta et al., 2014). Stakeholders also identified pollution and disease as perceived pressures. These pressures may interact with environmental stressors through effects on physiological and immune function, although such interactions were not tested here.

The accounts are consistent with exposure to multiple pressures, but the study does not establish their causal interactions. This distinction supports considering ecological pressures alongside governance capacity, rather than treating the interviews as evidence of specific recovery thresholds. A policy option is to investigate precautionary monitoring and adaptive responses that could reduce harvesting pressure during high-risk periods.

### 4.2. Governance fragmentation and uneven rules-in-use

Despite strong convergence on ecological drivers, respondents were often pessimistic about governance and management performance. The recurrent explanation was not an absence of rules, but a mismatch between formal regulations and the spatially heterogeneous, multi-gear and multi-use realities of nearshore clam grounds. This is consistent with the “*limits of governability*” perspective: governance effectiveness depends not only on rule content, but also on institutional fit, capacity, and legitimacy in dynamic systems (Jentoft, 2007).

Decentralisation created a mosaic of regulatory frameworks, monitoring capacity, and enforcement approaches across AACC. While regional tailoring can support context-specific management, interviewees perceived that heterogeneity has produced unequal treatment of gears and ports, delays in updating measures, and weak coordination across adjacent grounds—constraints that become acute where multiple modalities exploit the same population but face different requirements for reporting, access, effort limits, or catch controls. In such conditions, inconsistent rules can generate perceived injustice, create incentives to shift effort toward less regulated modalities, and make stock-based approaches harder to implement credibly (Nielsen, 2003).

Respondents also characterised decision-making as overly top-down and bureaucratic. Even where management plans and follow-up committees exist, participation was often characterised as limited in influence over timing and design of measures, an important concern because legitimacy is central to compliance in SSF, where enforcement alone is rarely sufficient (Nielsen, 2003; Ostrom, 1990).

### 4.3. Rules-in-use, compliance, and the credibility of institutions

A related theme was the distinction between formal rules and “*rules-in-use*”—how regulations are interpreted, implemented, monitored, and sanctioned in practice. This distinction is foundational in commons governance: formal rules do not determine outcomes unless users view them as legitimate and feasible, and unless monitoring and sanctioning are credible (Dietz et al., 2002; Ostrom, 1990). Several interviewees contrasted earlier periods of stronger guild self-regulation with later phases of more centralised decision-making, highlighting the perceived importance of combining administrative authority with locally grounded organisational capacity.

In this framing, compliance problems are not reducible to individual behaviour; they may also reflect institutional credibility. Where fishers perceive uneven rule application across modalities, inconsistent enforcement, or limited procedural fairness, willingness to comply may decline (Nielsen, 2003). Conversely, where fisher organisations maintain credible internal rules and sanctions, compliance can improve because the institutional system lowers ambiguity, increases legitimacy, and reduces incentives for opportunistic behaviour (Basurto, 2008; Ostrom, 1990).

### 4.4. Monitoring, enforcement constraints, and informal practices

Stakeholders repeatedly described divergence between formal rules and practices on the ground, emphasising off-auction sales and poaching. These practices can undermine traceability, distort effort and landings statistics, and erode legitimacy by creating perceptions of unequal burden, particularly where compliant fishers face sanctions while illegal extraction persists.

A key insight is that the feasibility of monitoring and enforcement is strongly modality dependent. Vessel-based operations return to predictable landing sites and pass through first-sale points, facilitating inspection. Shore-based fisheries—especially those targeting *D. trunculus* in very shallow waters—operate along beaches used intensively for recreation and can transport catches by car, reducing the effectiveness of port-based control strategies. Respondents also expressed concern that diffuse harvesting by beachgoers could contribute to unreported removals, conflict, and enforcement costs. The magnitude of these removals was not quantified by this study.

From a policy standpoint, informal practices should not be treated only as enforcement failures; they may also reflect institutional mismatch. Where transaction costs are high (e.g., compliance and depuration requirements), profitability is low, and rules are perceived as unevenly applied, incentives shift toward informal channels. Strengthening compliance, therefore, requires a dual approach: (i) enforcement designs suited to nearshore realities (e.g., mobile patrols and targeted seasonal controls on beaches) and (ii) legitimacy improvements that address distributional concerns and reduce contradictions across modalities (Nielsen, 2003; Ostrom, 1990).

### 4.5. Nearshore conflict and cross-sector coastal governance

Interviewees described conflicts with beach users, tourism priorities, and coastal works, underscoring that clam fisheries are embedded in crowded, multi-use nearshore landscapes. Fisheries conflict scholarship highlights that resource degradation and competing uses can intensify disputes, while institutional arrangements and stakeholder engagement play an important role in conflict resolution (Cánovas Molina & Gourguet, 2025).

The findings also resonate with broader concerns that SSF can be marginalised in coastal planning processes when their contributions are less visible or less politically salient, producing distributive conflicts over space and access (Bennett et al., 2015; Jentoft, 2007). Importantly, conflicts are not inherently “*abnormal*” in multi-use coastal systems; they may clarify values and power relations and create opportunities for dialogue and learning, provided that legitimate mechanisms for their articulation and resolution are available (Coser, 1956; Jentoft, 2007; Ostrom, 1990). Respondents’ accounts suggest such arenas exist more clearly for within-sector disputes (often mediated by guilds) but are weaker or absent for conflicts between fisheries and other coastal users, contributing to repeated, ad hoc resolution attempts and persistent feelings of marginalisation.

These dynamics imply that effective clam management cannot be confined to fisheries regulations alone. Aligning fishery needs with maritime spatial planning and coastal permitting—particularly around the timing and design of coastal works affecting sediment processes—requires clearer institutional coordination and structured opportunities for stakeholder dialogue to help prevent and resolve conflicts (Chang & Lin, 2016; Jentoft, 2007, 2017).

### 4.6. Implications for policy and management

The findings and the literature identify three policy options for consideration; their effectiveness was not evaluated in this study.

1. Mainstream climate-adaptive management under cumulative stressors. Interviewees identified high temperatures, storms, and rainfall variability as stressors, and the literature indicates that environmental variability can influence subsequent clam biomass, while temperature and salinity extremes can affect physiological condition and survival (Bento de Almeida et al., 2021; Soms-Molina et al., 2024). These considerations support integrating environmental indicators into management triggers, developing risk-based harvest control rules that can reduce effort rapidly during high-risk periods, consistent with the adaptive adjustment of fishing effort documented by Aguión et al. (2022), and implementing habitat-focused measures that protect sediment processes and avoid damaging coastal works during sensitive periods.
2. Increase coherence across gears and territories. Where different modalities exploit the same population, management objectives, data requirements, and effort controls should be harmonised to avoid perverse incentives and perceived injustice. Practically, this implies defining management units around stocks and fishing grounds, aligning monitoring protocols, and ensuring comparable catch accounting and traceability requirements across modalities if stock-based limits (e.g., TACs) are to be meaningful.
3. Institutionalise meaningful co-management and co-produced monitoring. LEK is well-suited to document ecological changes and identify locally appropriate management measures in dynamic nearshore contexts where scientific monitoring may be limited (Karnad, 2022). However, using LEK effectively also depends on selecting people with relevant knowledge and experience (Davis & Wagner, 2003).

The interviews highlight concerns about limited influence in decision-making. A policy implication is that participation should be substantive rather than merely procedural, with clearly defined roles for guilds and fisher organisations in rule design, participatory monitoring, and adaptive decision-making (Nielsen, 2003; Ostrom, 1990), paired with sustained administrative commitments to enforcement and cross-sector coordination.

### 4.7. Limitations and future research

As a LEK-based analysis, this study relies on perceptions and retrospective accounts that may be affected by recall bias and shifting baselines, and interview coverage was uneven across regions as some fisheries contracted. The design does not quantify the relative contribution of drivers or test causal pathways. The study also did not include an independent, standardised indicator of management effectiveness that was comparable across all study AACC. Consequently, assessments of management effectiveness represent stakeholder perceptions and should not be interpreted as externally validated measures of management performance. Future research would benefit from triangulating LEK with time-series analyses of landings and effort, spatially explicit habitat and coastal pressure indicators, and targeted monitoring of recruitment, size structure, and mortality events across modalities and regions.

Differences in interview conditions should also be considered when interpreting regional comparisons. Two interviews in Catalonia were conducted face-to-face, whereas all remaining interviews were conducted remotely. The interviewer was previously acquainted with four participants in Catalonia, but not with participants from the other study areas. These differences may have influenced rapport and the level of detail provided. Moreover, participants were recruited for their direct experience of clam fisheries rather than through probability sampling, and the sample should therefore not be considered statistically representative of regional fisher populations. Although basic demographic and professional characteristics were recorded, broader socioeconomic and cultural profiles were not systematically characterised, limiting our ability to assess their influence on responses. Regional comparisons should therefore be interpreted as patterns in stakeholder perceptions rather than as population-level differences.

Future comparative work could examine how coordination across fishing modalities, the feasibility of compliance, user participation, and integration with coastal planning relate to management outcomes. Such work would extend the institutional and spatial-planning perspectives discussed here (Chang & Lin, 2016; Jentoft, 2007; Ostrom, 1990).

### 4.8. Conclusions

By documenting perceived ecological change, examining institutional constraints and comparing stakeholder accounts across coastal settings, this study addresses the three objectives set out in the Introduction.

The accounts are consistent with H1 in identifying multiple pressures associated with clam decline, and with H2 in identifying perceived limitations in coordination, participation, rule implementation, monitoring and enforcement. Descriptive differences among study areas are also consistent with H3, although the study cannot separate contextual effects from differences in the composition of the interview sample. These findings do not establish causal interactions or independently measure management performance. They nevertheless identify practical priorities for coastal governance: clearer coordination across modalities, monitoring that draws on local knowledge, credible and feasible compliance arrangements, and decision-making that considers fisheries alongside other nearshore uses. The principal contribution is to show why accounts of ecological change and the institutions governing access and harvesting should be examined together when developing responses to declining clam fisheries.

## 4.9 Acknowledgments

This work was supported by Programa de Ayudas a Proyectos de I+D+I del Plan Complementario de Ciencias Marinas y del Plan de Recuperación, Transformación y Resiliencia de la Comunidad Autónoma de Andalucía, Project PCM_00122 to CR and MH. This study forms part of the ThinkInAzul programme supported by MCIN with funding from European Union NextGenerationEU (PCM_00122 and PRTR-C17.I1). We are grateful to the Departament d’Agricultura, Ramaderia i Pesca de la Generalitat de Catalunya, Dr. Jose Valencia from the Balearic Islands, the Conselleria d’Agricultura, Aigua, Ramaderia i Pesca de la Generalitat Valenciana, the Consejería de Agua, Agricultura, Ganadería y Pesca de la Región de Murcia, and the Consejería de Agricultura, Pesca, Agua y Desarrollo Rural (Junta de Andalucía) for providing the fisheries data. Finally, we greatly appreciate the helpful comments from the anonymous referees who evaluated the present work.

## Supplementary Material I

### Semi-structured Interview Protocol for the Analysis of Bivalve Fisheries along the Mediterranean coast of Spain and Atlantic Coasts of Andalusia

(English translation of the original Spanish-language interview protocol used for data collection.)

**Interview introduction:** The interview begins with a brief presentation of the project, an explanation of the interview’s objectives, and the verbal or written agreement to a confidentiality clause.

**General Information:**

- Name:
- Age:
- Stakeholder category:

⍰ Fisher
⍰ Fishers’ guild leader
⍰ Manager
- If fisher, specify fishing modality:

⍰ Hand-operated dredge (on foot)
⍰ Mechanised clam dredge
⍰ Towed dredge (“rastrell de cadenes”)
⍰ Towed dredge (“rastro”)
⍰ Hydraulic dredge
⍰ Other:
- Current status:

⍰ Active fisher
⍰ Former fisher
⍰ Other activity
- Main species targeted:

⍰ Donax trunculus
⍰ Chamelea gallina
⍰ Callista chione
⍰ Multiple
- Autonomous Community:

⍰ Catalonia
⍰ Valencian Community
⍰ Balearic Islands
⍰ Region of Murcia
⍰ Andalusia (Mediterranean)
⍰ Andalusia (Atlantic)
- Fishing zone or port:

**Note:** Gender was recorded by the interviewer based on visual observation, while years of fishing experience were obtained from information provided by participants during the interviews. Neither variable was included as a specific question in the original interview protocol. Retirement and changes to other occupations were distinguished based on participants’ explanations of why they had stopped fishing. These variables were assessed among the 70 fishers interviewed.

### Historical context and evolution of shellfishing

- Is there currently an extractive shellfishing activity in your Autonomous Community (AACC)? And in your specific fishing zone or port?
- Has there historically been shellfishing activity in the area / port?
- When did clam fishing begin in your AACC / zone / port?
- What species were or are targeted?
- What fishing gears were / are used (*e.g*., hand-operated dredges, mechanized clam dredges, towed dredges, or hydraulic dredge)? Can you describe how they work?
- Has fishing gear evolved or improved over time? How?
- Have any species or fishing methods disappeared? Which ones and why?
- How many fishers and vessels were historically involved in each fishing modality? (disaggregate). How many crew members per vessel? And now?
- Which zones or ports were the most productive? Are they still productive?
- When were zones or fisheries closed? By species?
- Why were they closed?
- Was the decline in catches gradual or sudden? Was it the same for all species? Did it occur simultaneously along the coast?
- What do you think were the main causes of the decline or collapse? Which were most important?
- What management measures were applied before the decline or collapse? Were extraordinary management measures introduced when catches began to decline? Which ones?
- How many fishers were affected?
- What did they do afterwards? Did they remain fishing? Switch activities? Receive financial or institutional support?

### Licenses, fishing zones, and effort

- Are / were there specific licenses for each modality or species?
- Can fishers target more than one species with the same license? On the same day?
- What criteria are/were used to determine the number of licenses?
- Are / were fishing zones clearly defined? Could fishers change zones?
- Are / were there multiple ports operating within the same fishing area?
- Are / were different modalities targeting the same species in the same area?
- If so, did this generate competition or conflicts between modalities?
- Were there differences in rules, access, or restrictions between modalities targeting the same species?
- Has this changed over time? How?
- Has fishing effort (number of fishers, vessels, or activity) increased or decreased over time? Why?
- Do you think fishing effort has been effectively controlled? Why or why not?

### Normative management and planning

- Who defines / defined management measures: administration, fishers, or both?
- To what extent are / were fishers involved in decision-making processes?
- What measures are / were applied (schedules, minimum sizes, quotas, fishing zones, mesh size, vessel engines, sales points)?
- Are / were these measures uniform across zones or specific species and areas?
- Do you think management measures are / were adapted to the specific characteristics of each fishery?
- Which measures were effective or ineffective? Why?
- Were extraordinary measures applied when catches declined? Which ones? Did they work? What failed? How did fishers react?
- What do you think about different regulations across neighboring regions for the same species? (*e.g*., minimum sizes). Would unified management be preferable?
- Do differences in regulations between regions create problems for fishers or for resource sustainability? How?
- Are there differences between official rules and what happens in practice? (*e.g*., enforcement, compliance).

### Collective organization and governance

- How were / are fishers organized? (fishing guilds, cooperatives, by zone or modality?)
- Did / do they implement stricter internal rules than the official ones? (e.g., shorter schedules, higher sizes, lower quotas)
- Were these internal rules respected by all?
- Did / do fishers from different modalities participate in joint meetings for shared species?
- Were / are all modalities equally represented in these decision-making spaces?
- Who participated in meetings with the administration (guild leaders, fisher representatives)?
- Do you think these representatives accurately reflect the interests of all fishers?
- Are there groups or modalities that are underrepresented?
- Do fishers trust these organizations (guilds, cooperatives) and their role in management? Why or why not?
- Have these forms of organization helped to improve management outcomes? How?

### Resource monitoring and control

- Were / are scientific or technical monitoring activities conducted? By whom? (guilds, local / regional government, researchers).
- Did monitoring change when catches started declining?
- Are / were regular inspections carried out? By whom (SEPRONA, regional authorities)? What did they focus on (gear, species size, zones, sales)?
- Do you think monitoring and control are effective? Why or why not?
- Are there differences in enforcement between modalities or areas?

### Off-the-record sales and illegal harvesting /poaching

- Are / were there direct sales outside of the official auction system (e.g., to restaurants)? Estimated percentage? And poaching?
- How have these percentages evolved? (by decade: 1980-1990, 1990-2000, 2000-2010, 2010-2020, 2020-present)

**Note:** Retrospective percentage estimates refer to the proportion of total catches, distinguishing between poaching and sales outside the official auction system by licensed fishers (see Section 2.8 of the main manuscript and Table 2). The intervals in the original interview protocol were intended to represent successive decades (1980s, 1990s, 2000s and 2010s), followed by the period from 2020 to the time of the interviews (2024–2025). The same temporal interpretation was applied in Table 2.

- How is / was illegal fishing addressed?
- Is / was it professionalised or casual (tourists, beachgoers)? Where is it more prevalent?
- Which species are/were most affected?
- Do / did illegal fishers comply with minimum sizes?

### Sanctions

- Are / were fishers subject to graduated sanctions when breaking rules?
- Who imposes / imposed these sanctions (fishing guilds, administration, Guardia Civil (police))?
- What sanctions apply / applied to illegal fishers? Are / were they sufficient?

### Conflict and resolution mechanisms

- Are / were there conflicts between fishers or modalities? How were they resolved?
- Are / were there conflicts with other users (*e.g*., swimmers, tourists, recreational navigation)?
- Are / were fishing zones marked during summer to avoid conflicts?
- Was / is there a Marine Spatial Management Plan? Could / can it help maintain the shellfishery?

### Governance and multilevel coordination

- Overall, do you think the shellfishing activity is/was well managed? Why or why not?
- Is it easy or difficult to manage? Why?
- Overall, do you think governance has been effective? Why or why not?
- Are/were there regular meetings involving all stakeholders (*e.g*., local, regional, national, EU authorities, tourism, consumers, and other fishers)?

### Catches and fishing effort

- How have catches evolved by species and modality over the decades?
- How have fisher numbers, vessels, and effort changed?
- Which species were targeted in each decade?

## Supplementary Material II

**Table S1.** Distribution of interview participants by study area and actor role/fishing modality. Values represent the number of interviewees (n). Zero indicates that no participants belonged to the corresponding category in that study area. The total sample comprised 94 participants.

| Actor role /<br>fishing modality | Catalonia | Valencian<br>Community | Balearic<br>Islands | Murcia | Mediterranean<br>Andalusia | Atlantic<br>Andalusia | Total |
| --- | --- | --- | --- | --- | --- | --- | --- |
| Mechanised clam<br>dredges fishers | 8 | 13 | 2 | 2 | 11 | 0 | 36 |
| Hand-operated<br>dredges fishers | 6 | 2 | 0 | 0 | 1 | 8 | 17 |
| Towed dredges 1<br>fishers | 5 | 0 | 0 | 0 | 0 | 0 | 5 |
| Towed dredges 2<br>fishers | 0 | 0 | 0 | 0 | 0 | 2 | 2 |
| Hydraulic clam<br>dredges fishers | 0 | 0 | 0 | 0 | 0 | 10 | 10 |
| Fishers' guild<br>leaders | 4 | 6 | 1 | 1 | 4 | 3 | 19 |
| Regional<br>managers | 1 | 1 | 1 | 1 | 0 | 1 | 5 |
| **Total** | 24 | 22 | 4 | 4 | 16 | 24 | 94 |

**Note:** “Vessel-towed dredges 1” refers to “rastrell de cadenes”, used only in Catalonia; “vessel-towed dredges 2” refers to “vessel-based hydraulic dredges” used only in Atlantic Andalusia (Gulf of Cádiz).

**Table S2.**
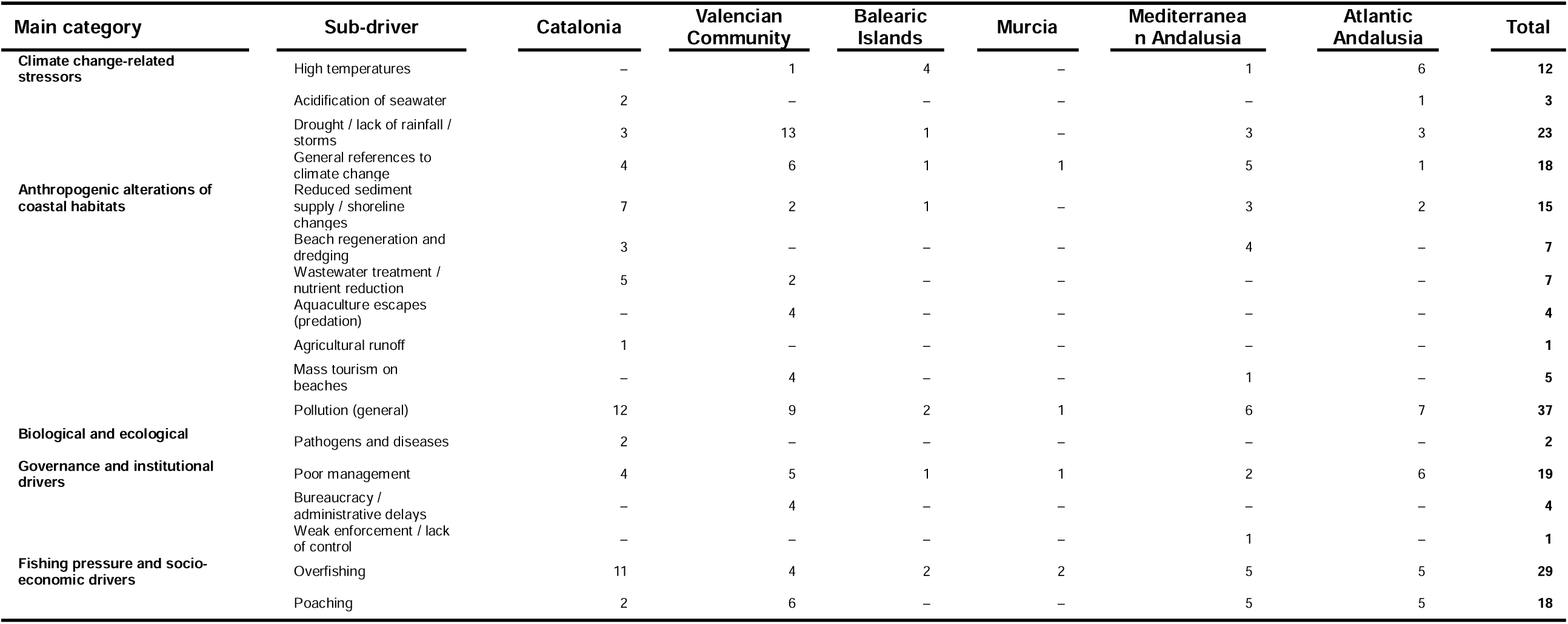
Frequency of references to specific stressor types mentioned by respondents across six study areas within five Autonomous Communities (AACC). **Each value represents the number of respondents mentioning each sub-driver within each AACC; repeated mentions of the same sub-driver by the same respondent were counted once. A respondent could be counted in more than one sub-driver when multiple perceived drivers were mentioned.** Dashes (–) indicate no mentions. Main categories correspond to those presented in Figure 3. Valid responses by AACC were: Catalonia (n = 22), Valencian Community (n = 22), Balearic Islands (n = 4), Murcia (n = 4), Mediterranean Andalusia (n = 15) and Atlantic Andalusia (n = 21), total 88 respondents.

